# Optimizing Single-Cell Long-Read Sequencing for Enhanced Isoform Detection in Pancreatic Islets

**DOI:** 10.1101/2025.04.30.651101

**Authors:** Maria S. Hansen, Christopher J. Hill, Lori Sussel, Kristen L. Wells

## Abstract

Alternative splicing is an essential mechanism for generating protein diversity by producing distinct isoforms from a single gene. Dysregulation of splicing that affects pancreatic function and immune tolerance has been linked to both type 1 and type 2 diabetes. Next-generation sequencing technologies, with their short read lengths, are limited in their ability to accurately detect splice variants. Long-read sequencing technologies offer the potential to overcome these limitations by providing full-length transcript information; however, their application in single-cell RNA sequencing has been hindered by technical challenges, including insufficient read lengths and higher error rates. Furthermore, cell types that produce high levels of a single transcript, such as islet endocrine cells, can obscure identification of lower abundance transcripts. In this study, we optimized a protocol for single-cell long-read sequencing in pancreatic islets to improve read length and transcript detection. Our findings demonstrate that 5’ library preparation protocols outperform 3’ protocols, resulting in better transcript identification. Furthermore, we show that targeted depletion of insulin transcripts enhances the detection of informative reads, highlighting the utility of transcript depletion strategies. This optimized protocol enables isoform-specific gene expression analysis and reveals differential transcript usage across the various cell types in pancreatic islets. By leveraging this approach, we gain deeper insights into the transcriptomic complexity and cellular heterogeneity within pancreatic islets.

**Article Highlights:**

- This study addresses the limitations of current single-cell long-read RNA-sequencing (sclrRNA-seq) technologies in detecting full-length transcripts and isoform diversity, particularly in pancreatic islets.
- We demonstrate that optimizing single-cell library preparation protocols reproducibly enhances read length and transcript identification in pancreatic islets.
- 5’ capture methods, combined with targeted insulin depletion and extended reverse transcription, significantly improved read length and isoform detection compared to standard protocols, while maximizing the number of informative reads.
- These improvements yield longer reads in single-cell experiments, substantially enhancing transcript identification and enabling more accurate analysis of isoform diversity.

## Introduction

Alternative splicing (AS) plays a critical role in generating protein diversity from the ∼22,000 known protein-coding genes, leading to the production of over 140,000 distinct transcripts [1]. This process allows for the generation of proteins with different amino acid sequences, impacting their functions and localization within the cell, and allowing them to respond readily to changes in the environment [2, 3]. Splicing dysregulation is a key factor in many diseases, including diabetes, either due to inherent mutations in splice sites or RNA-binding proteins (RBPs), or in response to changes in environmental conditions, such as inflammatory stress or hyperglycemia [4, 5]. In the context of type 1 diabetes (T1D), diversity in isoform expression has significant implications for pancreatic function and immune tolerance. For example, differential isoform expression of autoantigens IA-2 and G6pc2 between the pancreas and thymus has been proposed to contribute to the generation of autoreactive T cells in T1D [6, 7]. Furthermore, dysregulated splicing events have been observed in islets from individuals with type 2 diabetes (T2D), underscoring the importance of splicing regulation in maintaining proper cellular function and immune homeostasis [5]. As one specific example, SNAP-25, a component of the SNARE complex responsible for vesicle fusion and exocytosis, exists as two isoforms (SNAP-25a and SNAP-25b). In SNAP-25b-deficient mice, [Ca²⁺] elevations are prematurely activated and delayed in termination, and insulin secretion is increased [8].

Despite the critical need to detect splice variants in the context of diabetes, next generation sequencing (NGS) technologies remain insufficient for this task. Identifying isoform-specific gene expression requires sequencing reads that span multiple exons of the mRNA transcript. In the human genome, transcript lengths are estimated to average between 1,800 and 4,900 bp, with the mode of the distribution around 2,000 bp [9]. NGS technologies have read lengths of 150 base pairs, making it difficult to identify isoforms. In contrast, long-read sequencing technologies, such as PacBio and Oxford Nanopore Technologies, offer the generation of full-length reads that can capture the full RNA molecule, thereby providing a clearer picture of isoform diversity. Published single-cell long-read RNA sequencing (sclrRNA-seq) libraries often report shorter read lengths than expected, which may limit transcript coverage and isoform detection. For instance, a recent study reported a median read length of 900 bp for sclrRNA-seq of two cancer cell lines [10].

Advances in sequencing technologies, especially single-cell approaches, have revealed the complex heterogeneity within the pancreas, uncovering distinct functional and transcriptomic subpopulations across different cell types. In pancreatic islets, single-cell genomics and patch-seq have identified transcriptionally and functionally distinct beta-cell subpopulations directly linking gene expression to key physiological processes such as vesicle exocytosis [11]. This heterogeneity underscores the importance of characterizing splicing events and their resulting isoforms at the single-cell level. However, sclrRNA-seq technologies come with inherent limitations. Nanopore flow cells produce fewer reads than Illumina, with around 20,000 reads per cell for a 5,000-cell experiment, well below the typical 30,000–50,000 reads per cell common in NGS. Moreover, Nanopore’s higher error rate (1%) increases the likelihood of incorrect barcode and UMI assignments. To overcome these challenges, we have optimized a protocol for pancreatic islets that improves read length, advancing the utility of long-read sequencing for single-cell transcriptomics.

## Research Design and Methods

### Dissociation of pancreatic islets

10-week-old female C57BL/6 mice were obtained from Jackson laboratories. Pancreatic islets were isolated from mice under ketamine/xylazine/acepromazine anesthesia by collagenase delivery into the pancreas via injection into the bile duct. The collagenase-inflated pancreas was surgically removed and digested. After isolation, islets were dissociated using Accutase in a 37°C bead bath for 25-30 minutes. Single-cell suspension was filtered through a 40 mm filter and quenched in RPMI media + 10% FBS. Cells were washed again with RPMI +10% FBS and with PBS + 0.1% BSA. Single-cell suspensions were loaded into a Genomics Chromium targeting 4000 cells per sample.

### Dissociation of spleens

Spleens were isolated from mice under ketamine/xylazine/acepromazine anesthesia. Spleens were dissociated through a 70 µm strainer in cIMDM using a 3 mL syringe plunger. Cells were washed, centrifuged, and treated with 1 mL Ammonium-Chloride-Potassium (ACK) lysis buffer for 30 seconds, followed by dilution in cIMDM and a second spin. After one additional cIMDM wash, cells were resuspended in PBS + 0.1% BSA. Single-cell suspensions were loaded into a Genomics Chromium targeting 4000 cells per sample.

### scRNA-seq library preparation and insulin depletion

Single-cell libraries were prepared using either the Chromium Next GEM Single Cell 3ʹ Kit v3.1 or the Chromium Next GEM Single Cell 5’ Kit v2 following the protocol up to and including step 2.4, stopping just before fragmentation. For the optimized libraries, the following modifications were applied to the 5’ library preparation: 1 ul 10 mM dNTP solution (Thermo Scientific FERR0191) was added to the reaction in step 1.1. The extension time was increased from 45 minutes to 2 hours in step 1.5. 1 ul 10 mM dNTP solution was added to the reaction in step 2.2 and the extension time was increased from 1 minute to 3 minutes. Insulin depletion was performed on cDNA from step 2.4 of the 10X Genomics Chromium library preparation using the DepleteX^TM^ RNA Depletion Panel (Insulin) kit from Jumpcode Genomics. We followed the PacBio MAS-IsoSeq protocol (December 2022, Version 1.0) with the following modifications: During RNP Complex Formation (Step A), we used 0.9 ul Cas9 instead of 2.3 ul, and 1.6 ul Insulin Guide RNA instead of 4.0 ul Single Cell Boost Guide RNA. During Bead Cleanup (Step D), we used 50 ul (1X) AMPure XP Beads instead of 75 ul 1.5X SMRTbell Cleanup Beads.

Following insulin depletion, long-read libraries were prepared from the cDNA using Sequencing Kit V14 (Nanopore SQK-LSK114) and the PCR Expansion (Nanopore EXP-PCA001). For 3’ libraries, the *Ligation sequencing V14 — single-cell transcriptomics with 3’ cDNA prepared using 10X Genomics on PromethION (SQK-LSK114)* protocol was used. For 5’ libraries, the *Ligation sequencing V14 - Single-cell transcriptomics with 5’ cDNA prepared using 10X Genomics on PromethION (SQK-LSK114)* protocol was used. Short Fragment Buffer (SFB) was used for library preparation instead of Long Fragment buffer (LFB). Library Beads (LIB) were used for the flow cell priming mix stead of Library Solution (LIS). Libraries were sequenced on R10.4.1 flow cells on either a PromethION 2 Solo (P2S) or PromethION 2 Integrated (P2i).

### Bulk RNA-seq library preparation

Islets were isolated from two male C57BL/6J mice as described above. RNA was purified using the Qiagen RNeasy micro kit (Qiagen Cat # 74004). Bulk RNA-seq libraries were prepared using KAPA mRNA Hyper Prep kit (Roche 8098123702), following the KAPA mRNA HyperPrep Kit protocol (KR1352 – v7.21). Insulin depletion was applied between step 8 and 9 in the KAPA library prep protocol.

### Single-cell RNA-seq preprocessing

Single-cell libraries were processed using the EPI2ME single-cell workflow v1.1.0 from Oxford Nanopore Technologies for the identification of cell and UMI barcodes (https://github.com/epi2me-labs/wf-single-cell) using the mm10-2020-A mouse reference provided by 10x Genomics. The kit name was either 3prime or 5prime depending on the 10x library preparation and we provided the kit version v3 and v1 for the 3’ and 5’ samples we prepared respectively. We provided the kit version that was indicated in the publications associated with each published dataset. All parameters can be found on github (https://github.com/CUAnschutzBDC/sc-islet-longread-analysis/blob/61d52b6ef94ff974a394a9335dd65d28b9e6360e/01_wf_single_cell/trial_parameters.tsv) The pipeline was altered to bypass the StringTie step to use a consistent GTF file from the mm10 genome. These jobs were executed using Singularity and Nextflow. Only the reads that had a sequence quality higher than 7 were input into this pipeline.

We manually created a gene count matrix from the BAM outputs that were produced by the EPI2ME pipeline, and then utilized the emptyDrops function from the DropletUtils package [12] to identify cell barcodes. Using these barcodes, we subset the gene and transcript count matrices from the single-cell workflow and calculated the read lengths of cells that were identified using the barcodes. Read lengths were calculated from the BAM files by looking at the length of the query sequences for all primary reads and were subset further by matching barcodes (CB), whether the read was tagged with a gene (GN), and whether the read was tagged with a transcript (TR). Finally, we examined the BAM files and calculated the number of reads that were tagged with a gene and a transcript, before creating the final output plots.

### Detailed analysis of the 5’ modified islet dataset

The filtered data for the 5’ modified islet dataset was input into Seurat (v4.1.3) [13–17], using R (v4.2.3). Quality control metrics were calculated using the perCellQCMetrics function from the Scuttle package (v1.8.4) [18]. We used the DoubletFinder package (v2.0.3) [19] to remove cells that were identified as doublets, before running PCA. We tested several resolutions of UMAP before generating final clustering.

Cell types were initially identified using a reference dataset [20] and further refined based on the expression of known islet cell type markers. Dimensionality reduction and clustering was first performed using gene-level expression data, followed by a second round of clustering based on transcript-level expression. To evaluate the consistency between gene- and transcript-based cell type annotations, we calculated concordance between the two methods and visualized the results as confusion matrices using the Matrix (v1.5-4) [21] and pheatmap (v1.0.12) packages. Clusters identified by transcript-level expression were used for downstream analysis. To assess differential splicing, we conducted pseudobulk differential transcript usage (DTU) analysis using DTUrtle (v1.0.2) [22] alongside pseudobulk differential gene expression (DGE) analysis. Pseudobulk count matrices were generated by aggregating counts per replicate and cell type using Seurat. Differential expression was tested at the gene level using DESeq2 (v1.38.3), and differential transcript usage was tested at the transcript level using DTUrtle with sparseDRIMSeq (v0.1.2) for model fitting. Batch effects across replicates were corrected using ComBat-seq from the sva package (v3.50.0). Most figures were generated using the dplyr (v1.1.0), ggplot2 (v3.4.1), readr (v2.1.4), tidyverse (v2.0.0), MetBrewer (v0.2.0), and scAnalysisR (v0.0.0.9000) [23] packages. Further details of this pipeline can be found at our code repository:

https://github.com/CUAnschutzBDC/maria_long_read_081224/

Docker images can be found here:

RNA-seq preprocessing (STAR, Fastqc, cutadapt, featurecounts):

https://hub.docker.com/r/kwellswrasman/rnaseq_general

scRNA-seq (DTurtle, scAnalysisR, Seurat):

https://hub.docker.com/r/kwellswrasman/sclrseq_methods_r_docker

### Bulk RNA-seq analysis

Raw reads were trimmed using cutadapt (v4.8) and python (v3.10.14) [24] and aligned to the mouse GRCm38 genome using STAR (v2.7.11b) [25]. Gene expression was quantified with featureCounts (v2.0.6) [26] and differential expression analysis was conducted using DESeq2 (v1.44.0) [27] with R version 4.4.1. All plots were made using ggplot2 (v3.5.1) in R.

### NGS coverage

The 10k Human DTC Melanoma, Chromium GEM-X Single Cell 5’ and 10k Human DTC Melanoma, Chromium GEM-X Single Cell 3’ aligned bam files were downloaded from 10x genomics (https://www.10xgenomics.com/datasets). Coverage plots were generated using NGS plot (v2.63) [28].

### Transcript Coverage

To visualize the transcript coverage for each of our samples, we utilized the coverage function from the GenomicFeatures (v1.60.0) package. The reads output from our single-cell post-processing pipeline were aligned to a transcriptome reference generated from the GRCm38 Ensembl GTF, and subset based on matching barcodes and unique UMIs from our processed Seurat objects. For each sample, a coverage matrix of counts per transcript position was created for every transcript in the GTF file, then each normalized to be a length of 100. The counts were then normalized by read depth, and then again by applying a softmax transformation to generate matrix values that summed to 1 across each transcript, before a column mean provided a value per each 1-100 position per sample. These coverages were subset based on the number of exons per transcript, supplied by the GTF file, as well as the length of the transcripts, with a prerequisite that the transcript have >1 exon.

### Gviz plots

For visualizing our transcriptome-aligned reads on the genome, we utilized Gviz (v1.52.0) and the bam files output from our single-cell post-processing pipeline. Briefly, reads for each sample were aligned to a transcriptome reference generated using the GRCm38 Ensembl GTF file, and subsequently subset based on matching barcodes and unique UMIs from our processed Seurat objects. The final output bam files were generated by subsetting the genome-aligned bam files to only those reads that uniquely mapped to one transcript in our transcriptome alignment. These bams were then merged by replicate. For each of these merged bam files, an alignments track was created utilizing Gviz and displayed alongside the GRCm38 transcripts.

### Overrepresentation analysis (ORA)

Differential transcript usage (DTU) genes were subjected to KEGG pathway overrepresentation analysis (ORA) using gprofiler2 (v0.2.1) [29] with a custom background of all expressed genes. KEGG pathways were extracted from the enrichment results and ranked by p-value. The top 25 pathways were visualized as bubble plots.

### 3’ versus 5’ bias plot

Isoform-defining splice junctions were identified across datasets. Transcripts were classified based on whether their unique junctions fell in the 3′ region (last 25% of the transcript), 5′ region (first 25% of the transcript), central (middle 50% of the transcript) or both 5’ and 3’, and binned accordingly

## Data availability

All data will be made available on GEO upon publication.

### Code Availability

All source data supporting the findings of this study are provided. Custom analysis pipelines are available on GitHub at:

https://github.com/CUAnschutzBDC/sc-islet-longread-analysis.

## Results

### Evaluation of read lengths and isoform detection in published single-cell long read datasets

We aimed to identify isoform differences between islet cell types and subtypes using single-cell RNA-sequencing. To evaluate the ability of single-cell sequencing technologies to generate full-length reads, we reanalyzed previously published sclrRNA-seq datasets generated using 10x Genomics and Oxford Nanopore Technologies, focusing on their ability to capture full-length transcripts and detect isoform-specific transcript expression. Our analysis included seven sclrRNA-seq libraries from five different studies [10, 30–33]. The reanalysis revealed an average read length of 794 bp, and an average mode of 582 bp, compared to the expected mode distribution of ∼2,000 bp in the human genome [9] (Figure 1A). This discrepancy between the average read length and the expected transcript length underscores the ongoing challenge of capturing full-length transcripts. This shortfall in read length is important because it limits the transcript detection ability. Where gene detection ranges from 60-75% of total reads, transcript detection ranges from 30-60% of total reads (Figure 1B). These findings highlight the limitations of current sclrRNA-seq technologies in achieving comprehensive transcript-level resolution.

**Figure 1.**
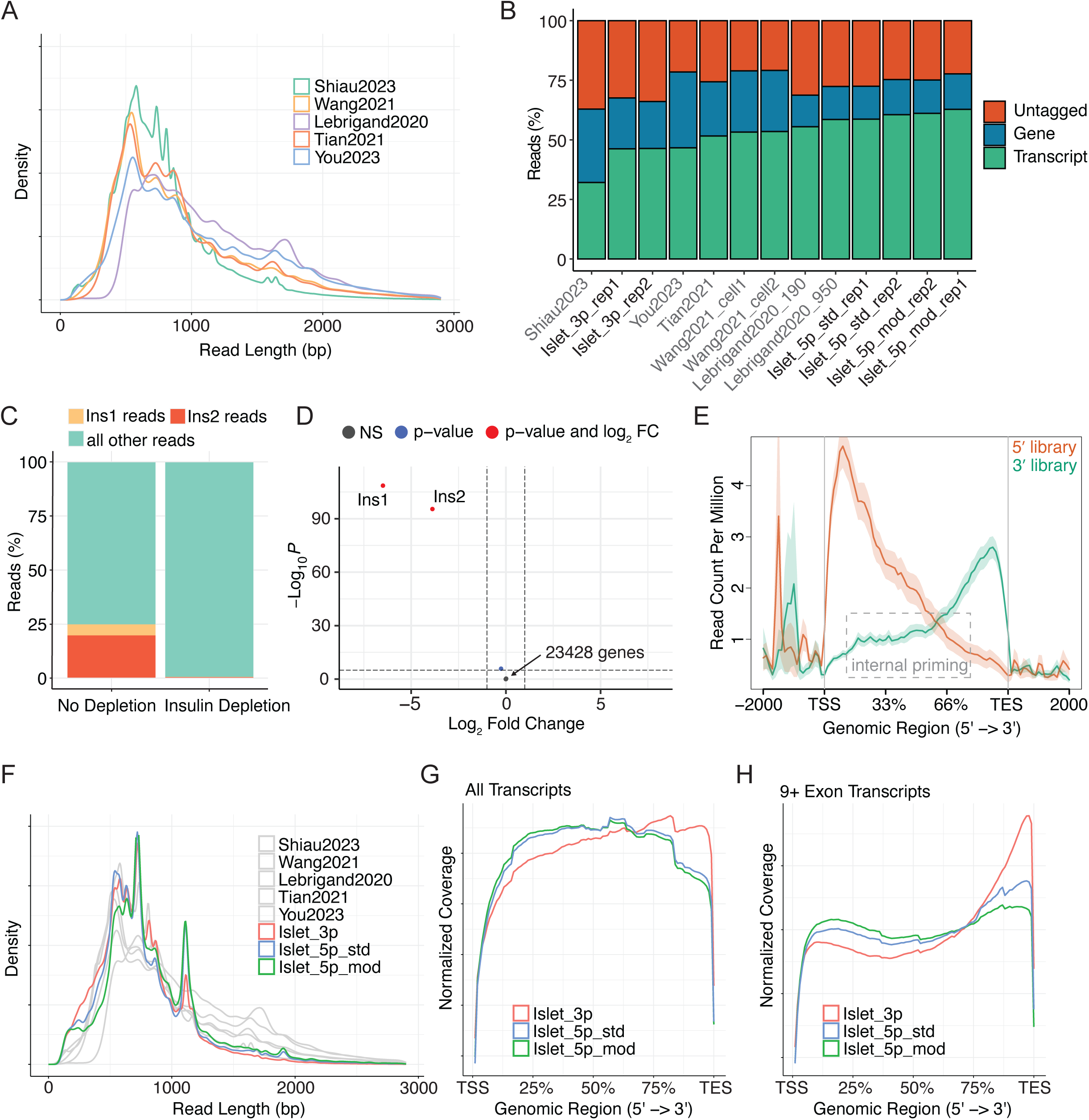
Read length and transcript identification comparison between single-cell long-read RNA-sequencing (sclrRNA-seq) libraries. (A) Read length distribution of published sclrRNA-seq libraries prepared with 10x Genomics and Nanopore technology. Biological replicates are included for datasets from Lebrigand et al., 2020 and Wang et al., 2021; other datasets are shown as single samples. (B) Proportion of reads across datasets where the gene is identified, the transcript is identified, or neither is identified. Shown are published reanalyzed datasets and six mouse pancreatic islet samples: two replicates prepared with 3′ 10x Genomics technology, two replicates with 5′ 10x Genomics technology, and two replicates with 5′ modified 10x Genomics technology (incorporating library preparation optimizations). (C) Proportion of reads aligned to *Ins1* or *Ins2* in a single-cell RNA-seq analysis of mouse pancreatic islets pre- and post-insulin depletion. (D) Volcano plot depicting differential gene expression between non-depleted and insulin-depleted bulk RNA-seq libraries from mouse pancreatic islets. A total of 23,431 genes is shown in the plot. (E) NGS coverage plot indicating read start sites across the genomic region. Libraries shown are 3′ and 5′ single-cell 10x Genomics preparations derived from human DTC melanoma cells. (F) Read length distribution comparison across mouse pancreatic islet sclrRNA-seq libraries, including two replicates prepared with 3′ 10x Genomics technology, two replicates with 5′ 10x Genomics technology, and two replicates with 5′ modified 10x Genomics technology. (G) Coverage plot showing Softmax-normalized read coverage across relative transcript positions for all multi-exon transcripts in mouse pancreatic islet scRNA-seq libraries prepared with 3′, 5′, or 5′ modified 10x Genomics technology. (H) Coverage plot showing Softmax-normalized read coverage across relative transcript positions for all transcripts with ≥9 exons in mouse pancreatic islet scRNA-seq libraries prepared with 3′, 5′, or 5′ modified 10x Genomics technology.

### Efficient and specific depletion of insulin from islet sequencing libraries generates enhanced read diversity

Analyzing transcript expression requires a higher overall read depth than gene expression analysis, as each gene is associated with multiple transcripts. Initial analysis of our sclrRNA-seq libraries of mouse pancreatic islets led to the discovery that the two mouse insulin genes, *Ins1* and *Ins2* made up 25% of the total reads, impeding our ability to achieve optimal read depth (Figure 1C). To overcome this issue, we incorporated an insulin depletion step into the protocol and validated the specificity and efficiency of the depletion in a bulk short-read RNA-sequencing library of mouse pancreatic islets. The depletion was remarkably efficient and highly specific: insulin transcripts were uniquely depleted, while all other genes remained unaffected (Figure 1D and Supplementary Table 1). The same insulin depletion was then applied to a single-cell pancreatic islet library followed by long-read Nanopore sequencing. Importantly, the insulin depletion was as efficient as in the bulk sample (Figure 1C). This strategy was applied to all subsequent pancreatic islet libraries generated for this study. Notably, the classification of beta cells does not rely on the presence of insulin transcripts. We demonstrate this by computationally removing insulin reads from a scRNA-seq library and repeating cell clustering (Supplementary Figure 1A-D).

### Protocol modifications enhance read length and transcript identification in islet single-cell long read libraries

Most high-throughput sclrRNA-seq methods rely on 10x Genomics single-cell capture and library preparation, which was originally optimized to generate and amplify shorter sequences, raising the question of whether it can effectively amplify full-length transcripts. 10x Genomics offers two types of transcriptomic profiling for single-cell RNA-seq: one that captures the 3’ end of transcripts and another that captures the 5’ end. Studies have shown that 3’ RNA libraries frequently contain internal priming artifacts [34] that would prevent the amplification of full length reads. To test for internal priming in 3’ vs 5’ libraries, we downloaded libraries generated using each technology in human melanoma samples from datasets created by 10x Genomics and analyzed the genomic coverage [28]. 3’ libraries exhibited a notably higher degree of internal priming compared to 5’ libraries, as evidenced by an increased number of reads mapping to the central regions of transcripts in genomic coverage plots (Figure 1E). Because reads generated through internal priming cannot span the full length of a transcript, this phenomenon likely contributes to the shorter read lengths observed in these libraries. To test this, sclrRNA-seq libraries were prepared from mouse pancreatic islets in parallel using both 3’ and 5’ capture technologies (n = 2 independent biological replicates per capture method). The libraries prepared with 5’ technology resulted in a subtle increase in longer reads, compared to the 3’ libraries (one-sided Wilcoxon test, p < 2 x 10^-16^) (Figure 1F and Supplementary Figure 2B). Furthermore, the 5’ library provided substantially improved transcript identification, increasing from 46.4 ± 0.1% to 59.7 ± 1.3% (Figure 1B).

To further improve the read length, several additional optimization steps were introduced into the islet 5’ library preparation protocol (Chromium Next GEM Single Cell 5’ Reagent Kits v2), including increasing the extension time from 45 minutes to 2 hours during GEM-RT Incubation and from 1 to 3 minutes during cDNA amplification, based on the approach outlined by Lebrigand et al. [30] and increasing the amount of dNTPs. These modifications resulted in longer reads than those from the 5’ library without modifications (one-sided Wilcoxon test, p < 2 x 10^-16^) (Figure 1F and Supplementary Figure 2B) and enabled far better transcript identification than any of the published datasets (62.0 ± 1.2%; Figure 1B). Additionally, the 5’ modified libraries showed improved transcript coverage (Figure 1G and Supplementary Figure 2C), with the most pronounced improvement observed in longer transcripts (Figure 1H and Supplementary Figure 2D). Overall, this emphasizes the preference for 5’ capture over 3’ capture and highlights the necessity for library prep optimizations to enhance the amplification of full-length reads.

Isolating high-quality RNA from pancreatic islets is notoriously difficult, primarily due to the presence of digestive enzymes, including RNases that are secreted by the exocrine pancreas. To explore whether a different cell type might yield still longer reads, we applied 3’, 5’, and 5’ optimized library preparations, as described above, to lymphocytes isolated from dissociated mouse spleens (n = 2 independent biological replicates per capture method). The 5’ lymphocyte samples demonstrated better transcript identification compared to the 3’ pancreatic islet samples (Supplementary Figure 2A). However, the 10x Genomics Chromium library prep modifications for the 5’ samples did not yield the same improvements in the lymphocyte samples as observed with the pancreatic islet samples (Supplementary Figure 2A, E-F). This suggests that the benefits of these optimizations might be tissue-specific and highlights the need for further refinements tailored to different tissue types.

### Isoform variants identified between alpha and beta cells and within beta cell subpopulations

With the improved library preparation, the optimized 5’ sclrRNA-seq datasets from mouse pancreatic islets were used to explore whether splicing changes could be detected from different cell types and cell states. Importantly, the 5’ modified sclrRNA-seq dataset, merged across replicates, allowed clear identification of all expected cell populations (Figure 2A-B and Supplementary Table 2). Importantly, despite depletion ∼95% of insulin transcripts, enough transcripts remain to identify insulin-expressing cells (Figure 2B). Furthermore, the analysis revealed that cell type identification remains robust whether using gene-level or transcript-level expression data for dimensionality reduction and clustering, with over 90% concordance between the two approaches (Figure 2D-G). This stability in broad cell type classification aligns with the understanding that major cell types are defined by distinct gene expression patterns. However, when examining substructure within these cell types, substantial differences emerged between gene-level and transcript-level analyses, with consistency ranging from 43% to 90% across subclusters (Figure 2I, Supplementary Figure 3). These findings suggest that while gene-level expression is sufficient for identifying major cell types, transcript-level analysis provides crucial insights into subtle variations within cell populations. Such variations may reflect different cell states, functions, or responses that are not captured by gene-level analysis alone.

**Figure 2.**
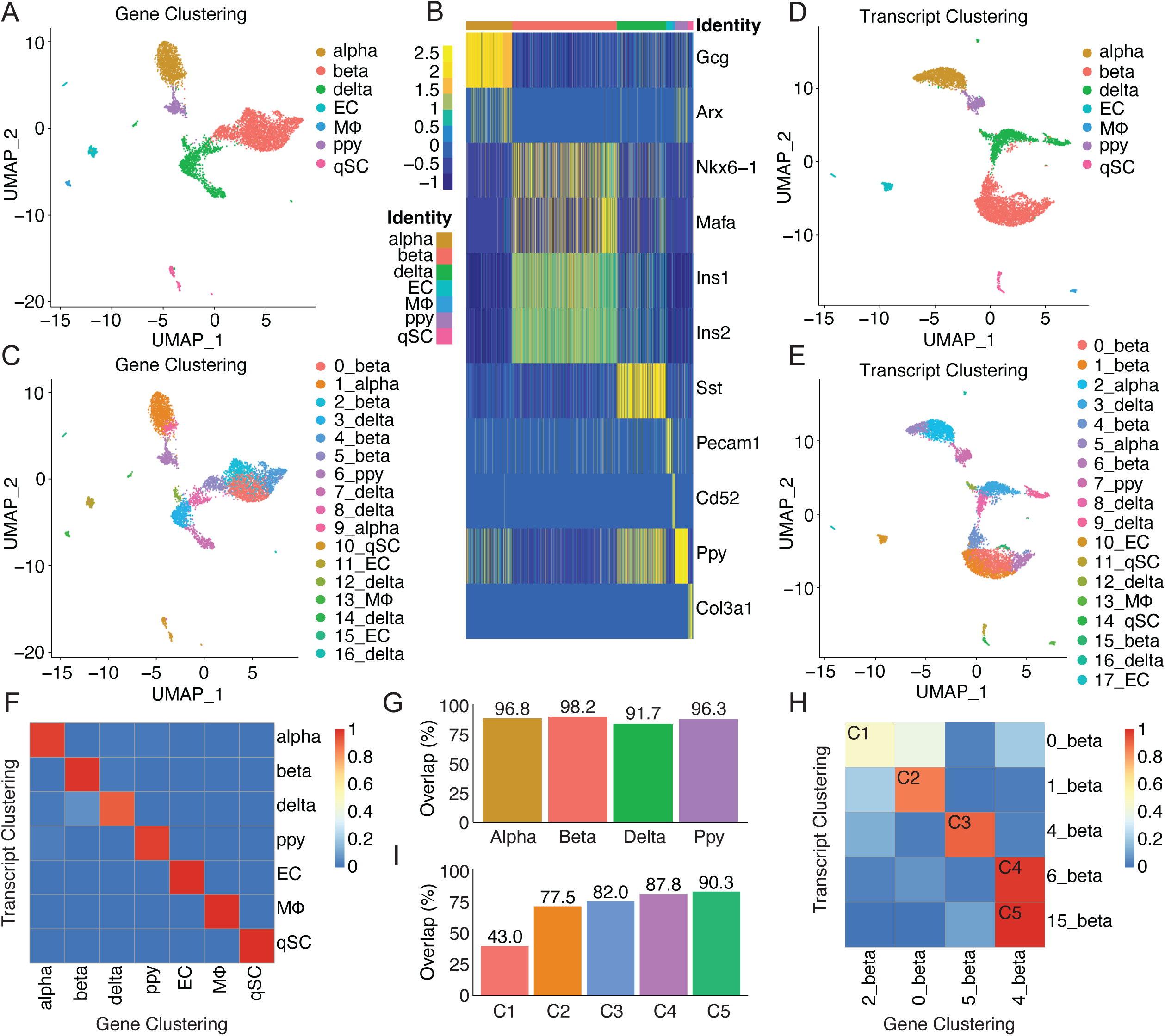
Comparison of single-cell clustering based on gene expression versus transcript-level expression. (A) UMAP projection of single cells from insulin-depleted mouse pancreatic islet sclrRNA-seq libraries prepared with 5′ modified 10x Genomics technology (two biological replicates merged), based on gene-level expression. Cells are colored by gene expression profiles reflecting major pancreatic cell types. (B) Heatmap showing expression of cell type-specific markers across single cells insulin-depleted 5′ modified libraries (two biological replicates merged), based on gene-level clustering. (C) UMAP projection of single cells based on gene-level expression, colored by gene expression profiles reflecting cell subpopulations. (D) UMAP projection of single cells based on transcript-level (isoform) expression. Colored by transcript expression profiles reflecting major pancreatic cell types. (E) UMAP projection of single cells based on transcript-level expression, colored by transcript expression profiles reflecting cell subpopulations. (F) Confusion matrix showing concordance in cell type identification between gene-based and transcript-based clustering. (G) Bar plot quantifying cell type concordance between clustering methods. (H) Confusion matrix showing low concordance in beta cell subpopulation identification between clustering methods. (I) Bar plot quantifying beta cell subpopulation concordance between clustering methods. C1–C5 indicate comparison 1 through comparison 5.

The primary strength of sclrRNA-seq lies in its ability to capture cell-specific isoform expression. To assess differential splicing, we performed pseudobulk differential transcript usage (DTU) analysis on merged 5’ modified replicates, alongside pseudobulk differential gene expression (DGE) analysis. A pseudobulk approach was chosen because it treats the biological sample, rather than individual cells, as the unit of replication, reducing pseudo-replication and yielding more robust and reproducible results [35, 36]. DTU analysis [22] identifies proportional differences in the transcript composition of a gene, comparing how much each transcript contributes to the total gene expression across conditions. Using this analysis, 218 DTU events were identified between alpha and beta cells (Figure 3A). We also detected 2,095 DGE genes, of which 56 overlapped between the DTU and DGE analyses (Supplementary Table 3). As a representative example of beta cell subpopulation analysis, we focused on the comparison between the 1_beta and 4_beta subpopulations, where we identified 316 DGE genes and 9 DTU genes, with 2 genes overlapping between the two analyses (Supplementary Table 4). Overrepresentation analysis (ORA) of DTU genes revealed enrichment in pathways such as synaptic vesicle cycle, calcium reabsorption, and oxidative phosphorylation in both alpha versus beta cells and 1_beta versus 4_beta comparisons (Figure 3B and Supplementary Figure 4A). Specifically, when comparing alpha and beta cells, we identified isoform-specific differences in *Gnas*, a key regulator of cAMP signaling and insulin secretion (Figure 3C and Supplementary Figure 4B). Additional DTU genes of interest in this comparison include *Pcsk2*, which encodes a prohormone convertase required for insulin processing, as well as *Prdx2*, and *Calm2*, which contribute to antioxidant defense and calcium signaling, respectively. Analysis of two beta cell subpopulations (1_beta and 4_beta) revealed distinct isoform usage of *Ndufs2*, a central component of oxidative phosphorylation and ATP production (Figure 3D and Supplementary Figure 4C). This comparison also highlighted isoform differences in *Stxbp1*, a gene important for insulin granule exocytosis. To assess whether 3′ or 5′ library preparation introduces bias in detecting transcript isoforms, such as preferential capture of variation near the transcript ends, we examined transcript identification based on splice junctions (Supplementary Figure 4D). This analysis showed that there was limited 3′ or 5′ bias, with differences largely driven by a few highly expressed transcripts. Representative gene plots illustrate that 5′ libraries generally provide reads spanning more exon junctions (Supplementary Figure 5), whereas 3′ libraries better detect certain long transcripts with substantial 3′ end variation (Supplementary Figure 6) and may disproportionately enhance detection of some transcripts due to internal priming (Supplementary Figure 7).

**Figure 3.**
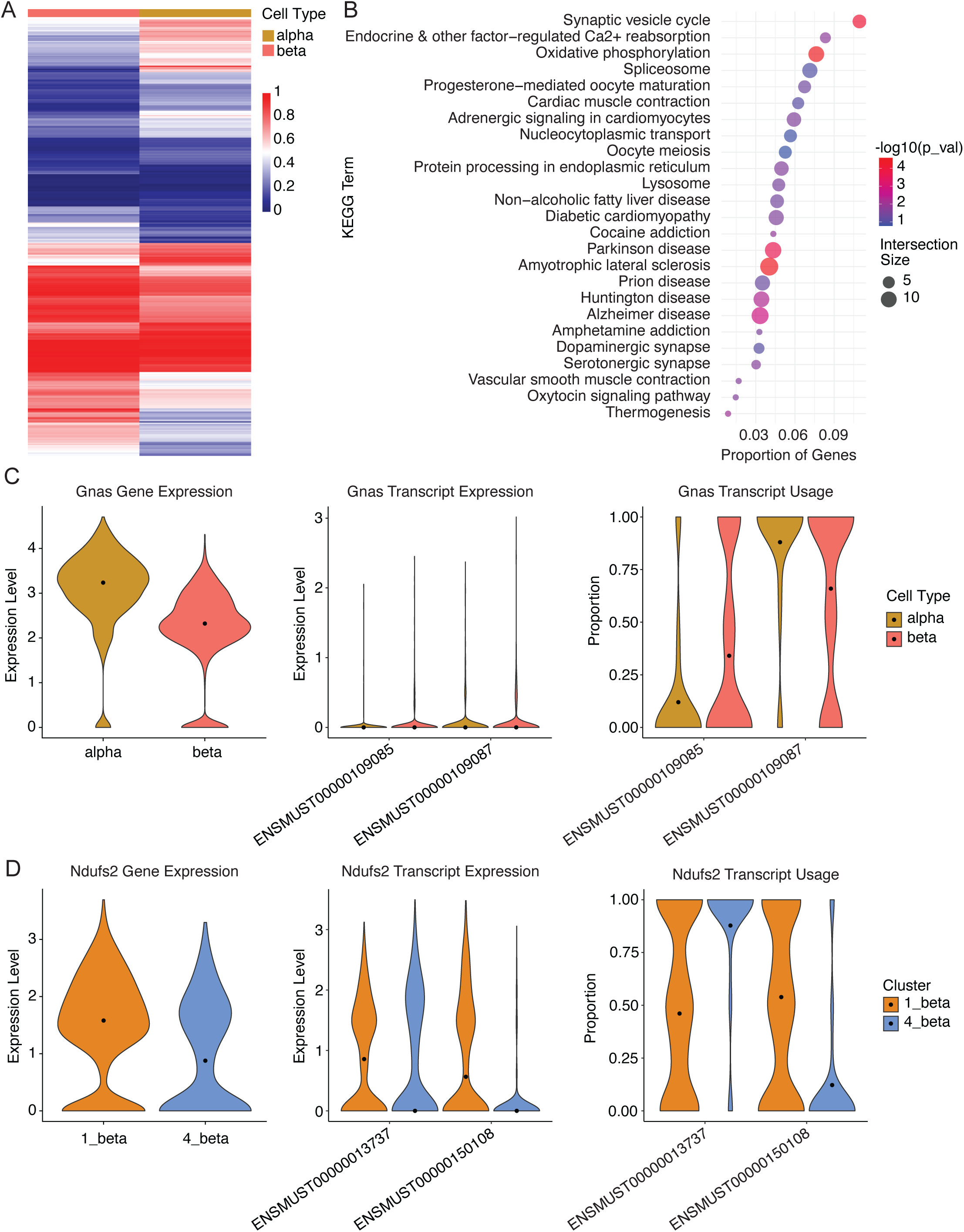
Differential transcript usage between cell types and cell subpopulations. (A) Heatmap showing differential transcript usage (DTU) between alpha and beta cells. Each row represents a transcript (significant at FDR < 0.05), and each column represents a cell type. Colors indicate the fraction of a gene’s expression contributed by each transcript within each cell type. DTU genes were identified from pseudobulk analysis of two biological replicates of mouse pancreatic islet sclrRNA-seq libraries prepared with 5′ modified 10x Genomics technology. (B) KEGG pathway overrepresentation analysis (ORA) for DTU genes between alpha and beta cells. The x-axis shows recall, i.e., the proportion of functionally annotated genes in the query that are associated with each pathway. Bubble size represents the number of DTU genes in the pathway, and bubble color indicates statistical significance (-log10 p-value). The top 25 enriched KEGG pathways are shown. (C) Differential gene expression (DGE), differential transcript expression (DTE), and differential transcript usage (DTU) analysis of *Gnas* between alpha and beta cells. DTU shows the relative contribution of each transcript to the gene’s overall expression. (C) DGE, DTE, DTU analysis of *Ndufs2* between beta cell subpopulations 1_beta and 4_beta.

## Discussion

This study demonstrates how an improved sclrRNA-seq library preparation protocol from isolated islets produces longer reads and increases the proportion of reads that can be confidently assigned to specific transcripts, improving the utility of long-read sequencing data for identifying splice variants and cellular heterogeneity in pancreatic endocrine cell populations. Specifically, this study demonstrates that islet sclrRNA-seq libraries prepared with 5’ protocols outperform those prepared with 3’ protocols. While neither library type showed a strong overall 3′ or 5′ bias, 3′ libraries are prone to internal priming, whereas 5′ reads tend to cover more exon junctions, providing more complete isoform identification. Enhancements to the 5’ library preparation further improves read length and transcript tagging efficiency in pancreatic islets. Furthermore, depleting insulin transcripts from the pancreatic islet libraries proved to be a highly effective strategy for maximizing informative reads, demonstrating the broader potential of targeted transcript depletion in single-cell RNA-sequencing experiments.

While the modified 5’ protocol significantly improved read length in islet samples, lymphocyte samples showed significant improvement only with the unmodified 5’ protocol, with no additional benefit from the modifications. This indicates that individual cell types will require unique modifications and further optimizations. Despite the significant improvements in read length achieved with the modified protocol, it did not meet expectations for full-length transcript coverage. Achieving this goal will require further modifications to the 10x chemistry, including adjustments to the master mix and reverse transcriptase.

Although full-length coverage was not achieved for all transcripts, we successfully analyzed transcript expression and identified differential transcript usage across cell types and cell subpopulations. These advancements are critical for uncovering the full complexity of transcriptomes and hold immense potential for broad application across tissues, enabling deeper insights into cellular heterogeneity, isoform regulation, and functional diversity. While scRNA-seq allows the identification of distinct cell populations and subpopulations, the precise number and composition of clusters can be influenced by analytical choices such as clustering resolution, preprocessing, and dimensionality reduction, supporting previous analyses showing that subpopulations may not be consistently defined across studies [37]. In addition, transcript capture and read depth can influence subpopulation detection: while adequate coverage suffices for gene-level conclusions, even greater depth is needed to resolve transcript-level differences, as each gene is often associated with multiple transcripts. Despite these limitations, single-cell transcriptomic analyses still provide valuable insights into isoform-level regulation within specific cell types. Investigating these variations at the single-cell level allows us to uncover the intricate heterogeneity within tissues, offering a deeper understanding of the functional and transcriptional diversity that would otherwise go unnoticed. Understanding splicing dysregulation in pancreatic islets is particularly important, as it may reveal how alternative splicing shapes beta cell function, immune tolerance, and beta cell susceptibility in diabetes.

## Acknowledgments

We thank Laura White, PhD, and Jay Hesselberth, PhD, for their guidance and support with Nanopore long-read sequencing technologies. We thank Mia Smith, PhD, for guidance on spleen sample preparation. We also thank Scott Beard, BDC Cytometer Core Manager, for islet and spleen isolations.

## Author Contributions

M.S.H. was responsible for data acquisition and prepared the original manuscript. K.L.W. and L.S. reviewed and edited the manuscript. C.J.H. and K.L.W. developed the computational pipelines. M.S.H., C.J.H., and K.L.W. contributed to data analysis and the graphical presentation of results. All authors contributed to the study’s methodology and conceptualization. K.L.W. is the guarantor of this work and had full access to all data in the study and takes responsibility for the integrity of the data and the accuracy of the data analysis.

---

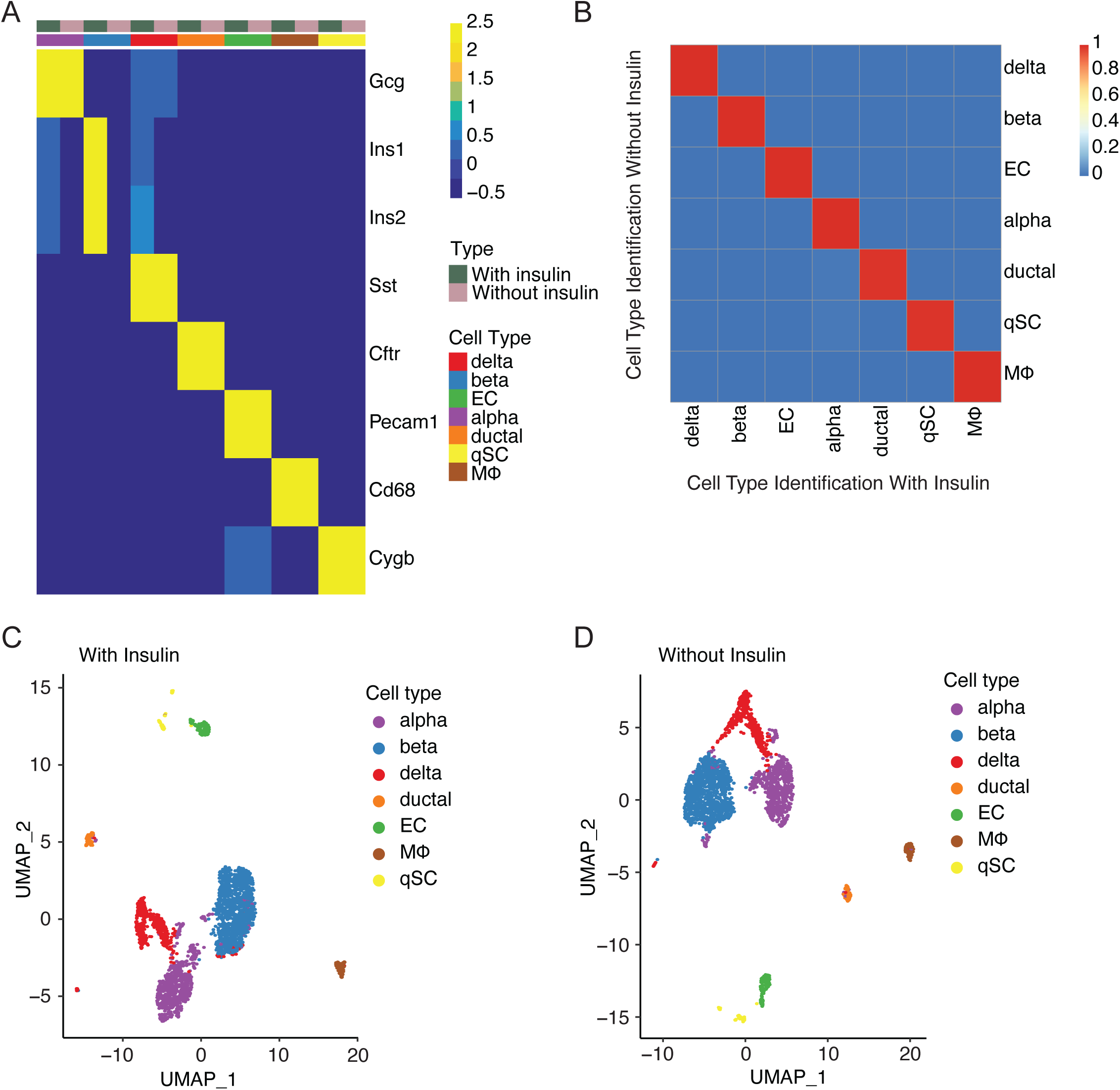
Cell type identification in mouse pancreatic islets with and without insulin transcripts. (A) Heatmap showing expression of cell type–specific markers across clusters defined by gene expression in a dataset including insulin compared to the same dataset from which insulin transcripts have been computationally removed. (B) Confusion matrix comparing cell type assignments between datasets with and without insulin transcripts. (C) UMAP projection of single cells from dataset with insulin, colored by cell type. (D) UMAP projection of single cells from dataset without insulin, colored by cell type.

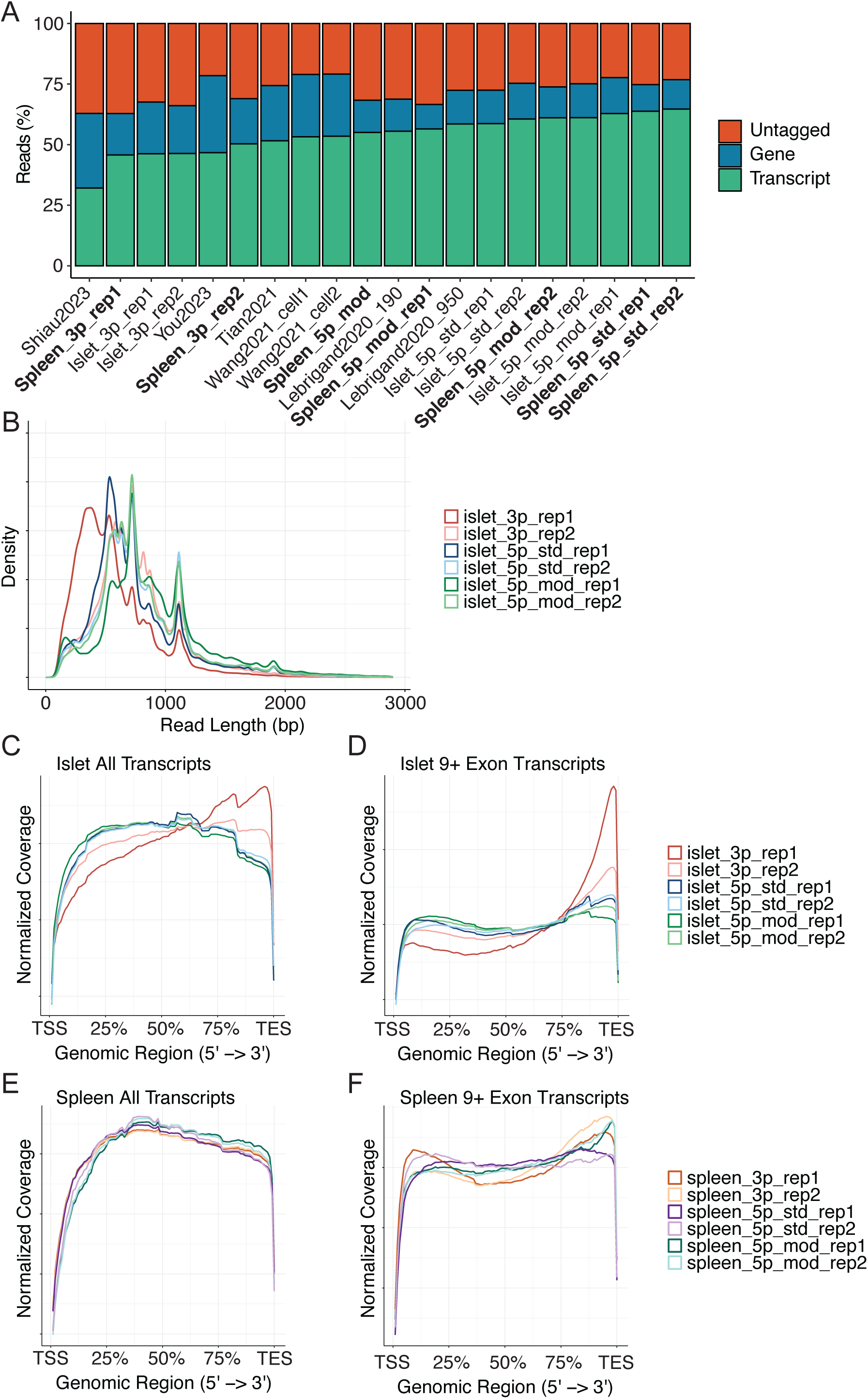
(A) Proportion of reads across datasets where the gene is identified, the transcript is identified, or neither is identified. Shown are six mouse pancreatic islet samples and six mouse spleen samples generated in this study, alongside published reanalyzed datasets. Each tissue from this study includes two biological replicates prepared with 3′ 10x Genomics technology, two with 5′ 10x Genomics technology, and two with 5′ modified 10x Genomics technology. (B) Read length distribution of mouse pancreatic islet scRNA-seq libraries, separated by replicate. Includes two replicates prepared with 3′ 10x Genomics technology, two replicates with 5′ 10x Genomics technology, and two replicates with 5′ 10x Genomics technology with library preparation optimizations. (C–F) Coverage plots showing Softmax-normalized read coverage across relative transcript positions in mouse pancreatic islet (C, D) and spleen (E, F) scRNA-seq libraries prepared with 3′, 5′, or 5′ modified 10x Genomics technology. Samples separated by replicate. Panels C and E include all multi-exon transcripts, while panels D and F include only transcripts with ≥9 exons.

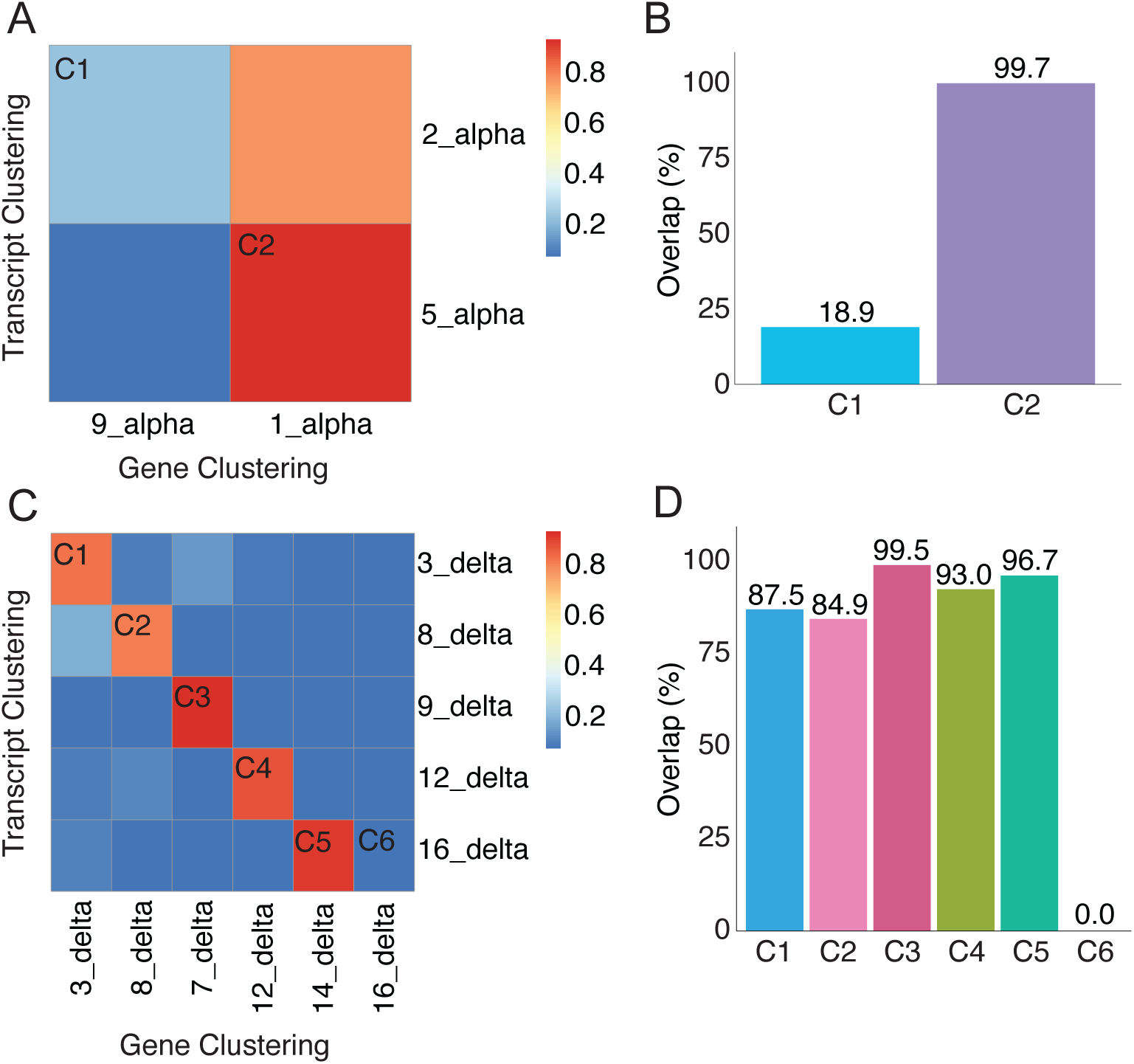
(A) Confusion matrix showing concordance in alpha cell subpopulation identification between gene-based and transcript-based clustering. (B) Bar plot showing concordance of alpha cell subpopulations across clustering methods. Concordance is quantified for two comparisons: C1 (comparison 1) and C2 (comparison 2). (C) Confusion matrix showing low concordance in delta cell subpopulation identification between clustering methods. (D) Bar plot quantifying delta cell subpopulation concordance between clustering methods quantified for comparisons C1-C6.

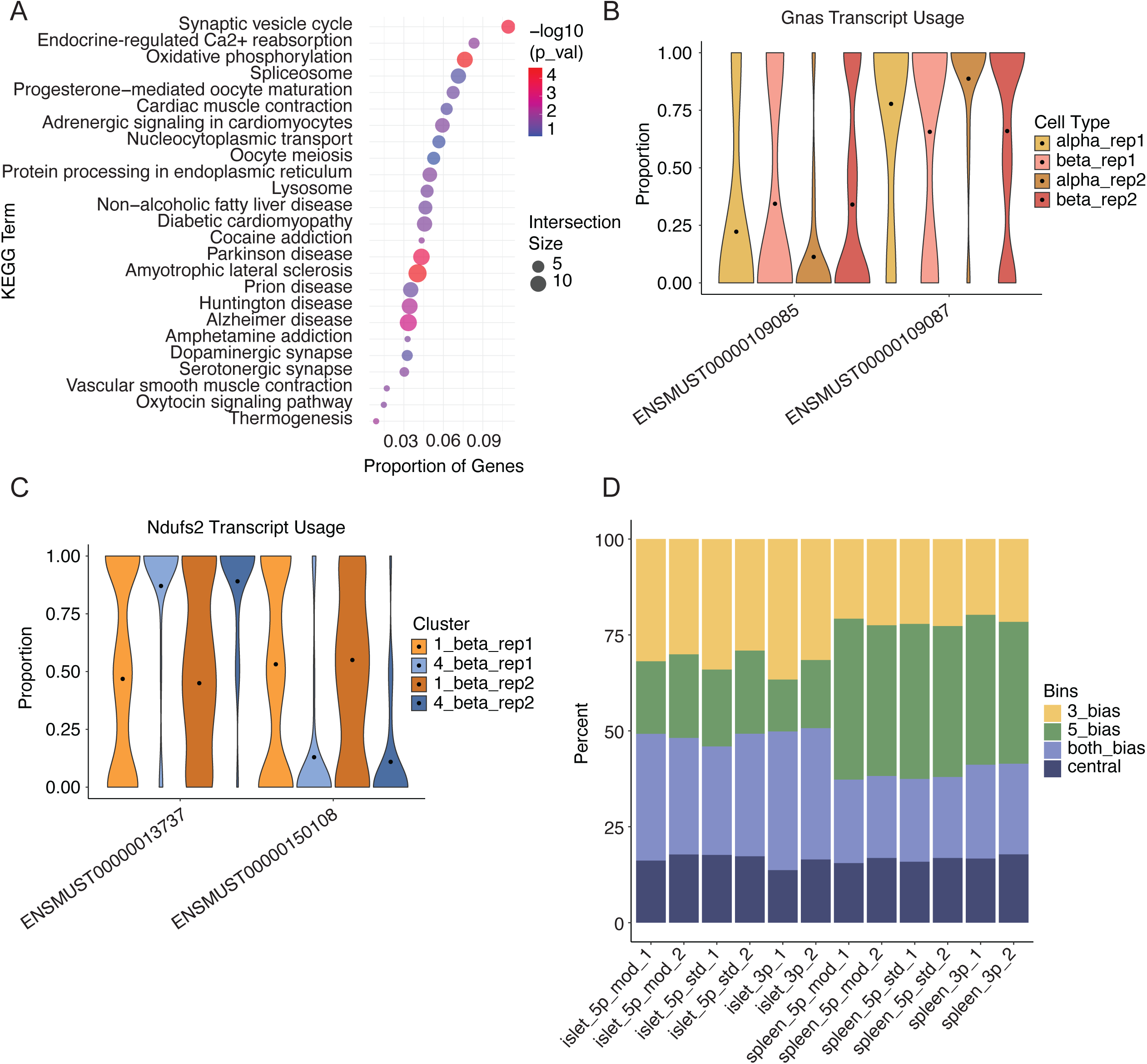
(A) KEGG pathway enrichment for DTU genes between beta cell subpopulations 1_beta and 4_beta. The x-axis shows the proportion of functionally annotated genes in the query that are associated with each pathway. Bubble size represents the number of DTU genes in the pathway, and bubble color indicates statistical significance (-log10 p-value). The top 25 enriched KEGG pathways are shown. (B) Differential transcript usage (DTU) analysis of Gnas between alpha and beta cells, separated by biological replicate. (C) DTU analysis of Ndufs2 between beta cell subpopulations 1_beta and 4_beta, separated by biological replicate. (D) Bar plot showing 3′ versus 5′ bias across libraries prepared with different technologies. Transcripts are classified by whether the junctions that uniquely identify each isoform fall in the 3′ region or 5′ region of the transcript, or both.

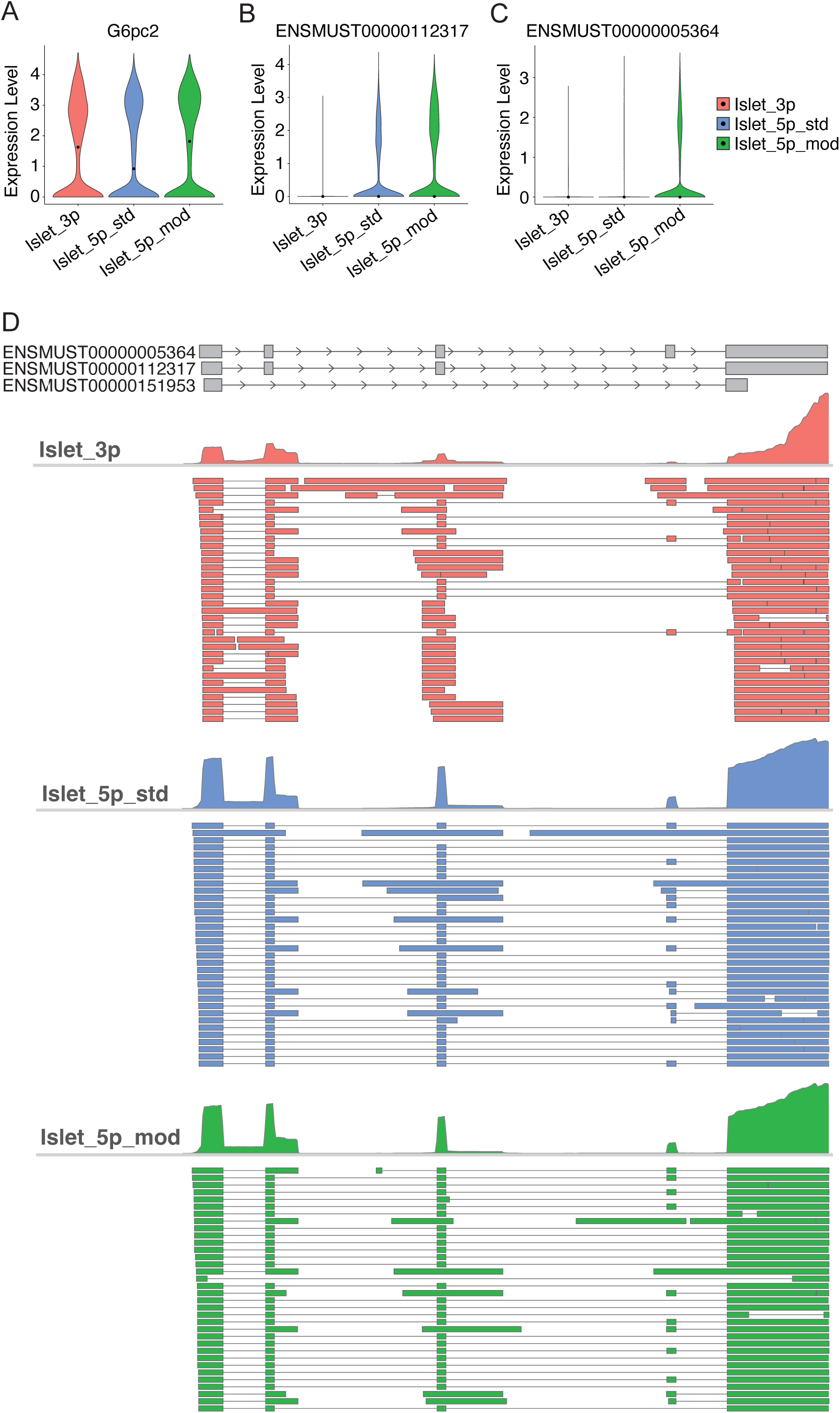
G6pc2 gene and transcript expression across mouse pancreatic islet libraries. (A) Gene-level expression of G6pc2 in 3′ modified, 5′ standard, and 5′ modified libraries (two biological replicates merged for each library). (B) Transcript-level expression of ENSMUST00000112317, isoform of G6pc2, in the same libraries. (C) Transcript-level expression of ENSMUST00000005364, isoform of G6pc2, in the same libraries. (D) Gviz plot showing a gene model track of select transcripts of G6pc2, and an alignments track including all reads uniquely aligned to any transcript of G6pc2.

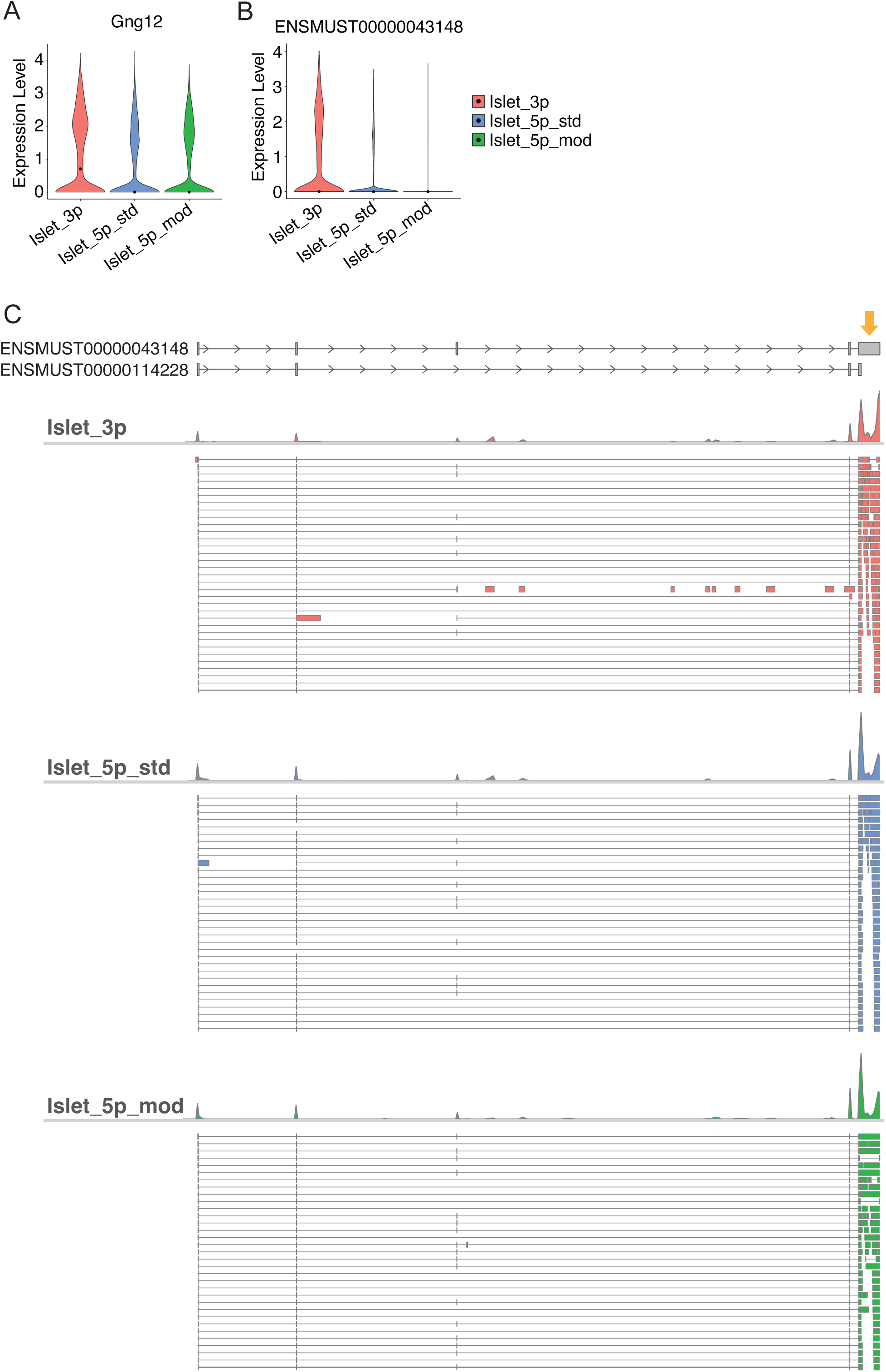
Gng12 gene and transcript expression across mouse pancreatic islet libraries. (A) Gene-level expression of Gng12 in 3′ modified, 5′ standard, and 5′ modified libraries (two biological replicates merged for each library). (B) Transcript-level expression of ENSMUST00000043148, isoform of Gng12, in the same libraries. (C) Gviz plot showing a gene model track of select transcripts of Gng12, and an alignments track including all reads uniquely aligned to any transcript of Gng12. Arrow indicates a long exon at the 3’ end of the transcript.

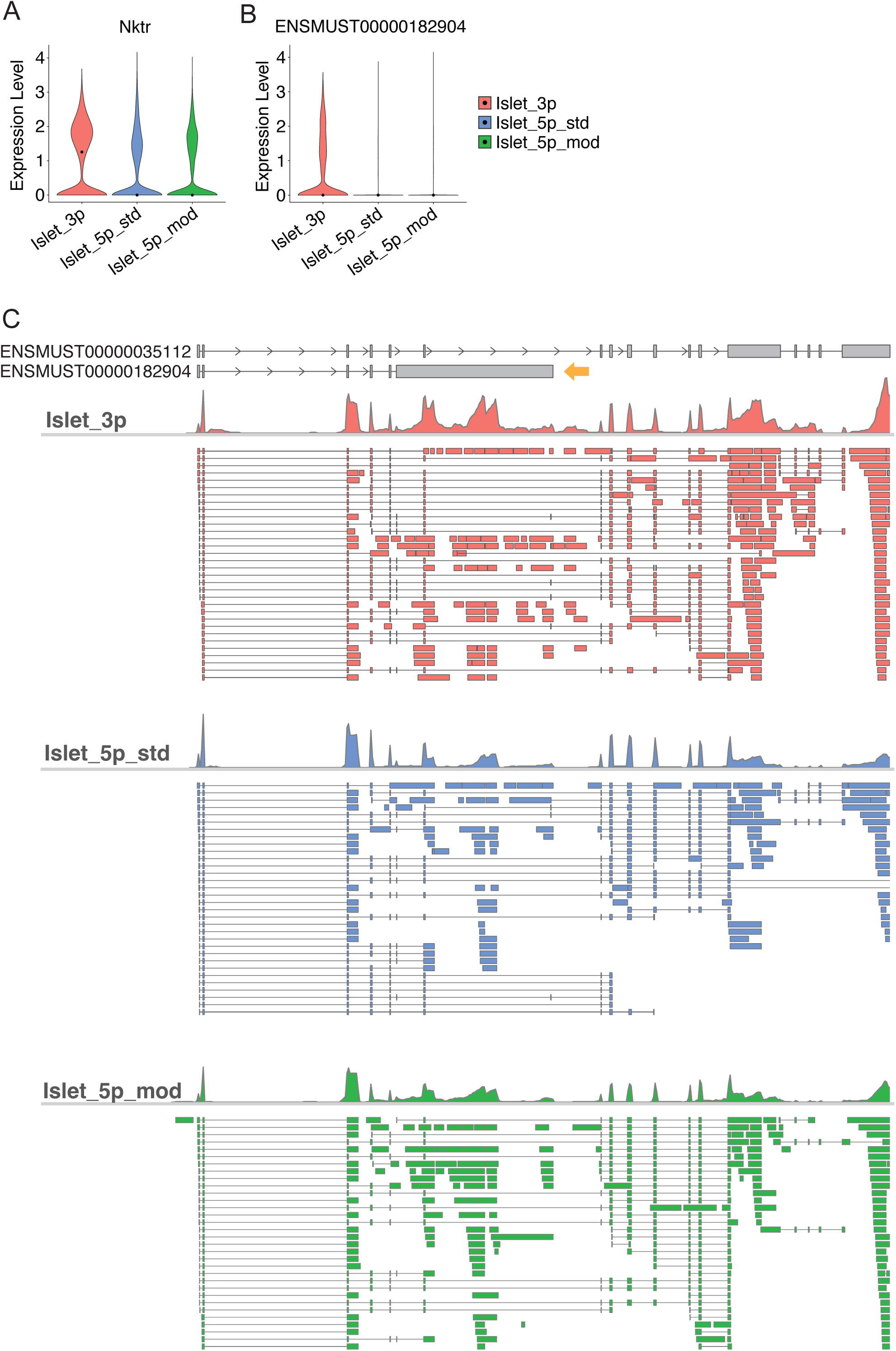
Nktr gene and transcript expression across mouse pancreatic islet libraries. (A) Gene-level expression of Nktr in 3′ modified, 5′ standard, and 5′ modified libraries (two biological replicates merged for each library). Transcript-level expression of ENSMUST00000182904, isoform of Nktr, in the same libraries. (C) Gviz plot showing a gene model track of select transcripts of Nktr, and an alignments track including all reads uniquely aligned to any transcript of Nktr. Arrow indicates retained intron captured by internal priming.

